# Constructing Predictive Cancer Systems Biology Models

**DOI:** 10.1101/360800

**Authors:** Jennifer A. Rohrs, Sahak Z. Makaryan, Stacey D. Finley

**Author notes:** J. A. Rohrs, S. Z. Makaryan, and S. D. Finley are with the University of Southern California, Los Angeles, CA 90089 USA (J.A.R.; S.Z.M.; S.D.F., corresponding author).

## Abstract

Systems biology combines computational modeling with quantitative experimental measurements to study complex biological processes. Here, we outline an approach for parameterizing and validating a systems biology model to yield predictive tool that can generate testable hypotheses and expand biological understanding.

## I. INTRODUCTION

When properly constructed and validated, computational systems biology models can provide unique insight into the complex biological processes that affect the growth, spread, and treatment of cancer. The computational models can be used to simulate many different experimental conditions, predict the effects of cancer drugs, evaluate alternative treatment protocols, and even identify new drug targets. Computational modeling can provide novel quantitative and mechanistic insight and reduce valuable resources invested in pre-clinical and clinical studies. However, development of predictive models is not trivial. It requires an iterative approach to identify an appropriate network structure and estimate the parameters such that the model can predict the outcome of different perturbations to the system. Here, we focus on parameterizing and validating mechanistic systems biology models and suggest how to troubleshoot issues that commonly arise.

## II. OVERVIEW OF METHODS

We describe an iterative approach of model construction (**Fig. 1**), with a focus on model optimization and validation. We present some of the lessons we have learned from training and validating cancer systems biology models.

**Figure 1:**
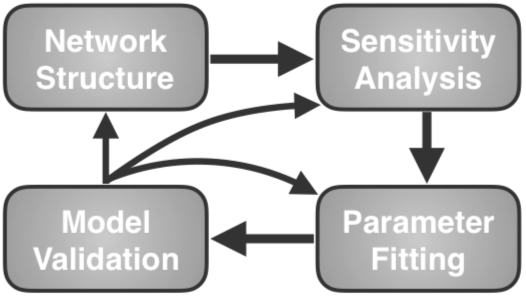
The model structure is specified and the values of influential parameters are estimated by fitting to experimental data. If model predictions do not match validation data, assumptions made in previous steps can be reevaluated to improve the predictive ability of the model.

### A. Network structure

Previous entries in this handbook have described the basic steps for building an ordinary differential equation (ODE) based mechanistic computational model relating to cancer, and provide a good starting point for model construction [1-2]. A mechanistic model requires a detailed understanding of the biological system being analyzed. After implementing the biology mathematically, it is important to consider the type and amount of data that is available to fit the model. Each entity measured needs to be represented as an output of the model. While quantitative data is preferred, qualitative observations can also be utilized. For example, if a system shows a transient response to one input and a sustained response to another, these behaviors can be enforced during parameter fitting. Importantly, some of the experimental data is used for training in the parameter fitting step, while a separate set of data is set aside for model validation.

Although the ODEs that govern species’ interactions can be written by hand, this process can be cumbersome and error-prone, particularly in the case of multiple binding or modification sites on a single protein. To circumvent this, a rule-based formalism, like BioNetGen [3], can be used. BioNetGen populates the ODEs for every species permutation possible, based on a minimal set of rules that define the molecular interactions. The output of this step is a set of ODEs that can be solved to simulate the dynamics of the biological system.

### B. Parameter sensitivity analysis

Frequently in computational models of biological systems, there are many more parameter values to specify than data points to which they can be fit. It is often the case that only a subset of the many parameters control the output, and it can be difficult to converge on a meaningful set of parameter values. Thus, it is important to identify the parameters that most significantly affect the model outputs. Parameters that are not influential can be held constant based on reasonable estimates from the literature, while influential parameters can be fit to the data.

A global sensitivity analysis, like the extended Fourier amplitude sensitivity test (eFAST) [4], can be used to quantify how sensitive the model output is to the parameter values. With eFAST, all of the parameters are varied at the same time, within specified ranges, and sampled at different frequencies. The Fourier transform of the model output can then be used to identify which frequencies, corresponding to individual parameters, contribute most to the changing output. This method calculates the first-order sensitivities for each parameter, a local sensitivity measurement, as well as the total sensitivity, which measures higher order interactions between multiple parameters.

Often, in biological systems, parameters can be correlated such that changes to one parameter are compensated for by proportionately changing another. This indicates that only one of the parameters can be determined, while the other is “non-identifiable”. To determine which parameters are identifiable, one can calculate the correlation between each pair of model parameters [5-7]. The output of the sensitivity and identifiability analyses is a set of identifiable parameters that significantly influence the model output, which will be fit to the experimental data.

### C. Parameter fitting

Parameter fitting can be both difficult and time consuming, particularly if there is a wide range over which the parameters can vary. We have implemented a two-tiered approach to address this issue. First, a global fitting algorithm, like particle swarm optimization (PSO), can be used to efficiently search a wide parameter space. PSO mimics the way that a swarm of bees searches for a new nesting place [8]. The algorithm begins with many different particles (parameter sets) scattered randomly throughout the parameter space. The model is run with each particle, and we calculate the error between the model predictions and the training experimental data. The particles then communicate their error to one other and move through the parameter space toward the lowest error, with some stochasticity to fully explore the range of parameter values. This process is repeated hundreds of times, ultimately arriving at the parameter set with the lowest error. We execute the whole PSO algorithm multiple times to generate a handful of sets that allow the model predictions to match the training data.

In the second step, the estimated parameter sets from the global fitting algorithm are fine-tuned using a local parameter fitting algorithm, such as nonlinear least-squares optimization to find the local minimum around the region identified by PSO. The output of this step is a series of parameter sets that can all fit the experimental training data well.

### D. Model validation

Model validation ensures that the model is able to predict data not used in the training process. Each set of parameters estimated in the parameter fitting step needs to be independently tested in the model to determine how well the model predictions match the validation data. Parameter sets that do not accurately reflect the validation data are discarded. Once a group of validated parameter sets is determined, it is also useful to analyze the median and standard deviation for the parameter values of that group. This can provide insight into how robust the model is and how heterogeneity in the model parameters may affect the output. If the model with the best fit parameters does not match the validation data, then the assumptions made in the previous steps need to be reevaluated. The model structure is an obvious step to return to; however, the initial starting values of the parameters and the fitting algorithm inputs may also need to be changed.

Finally, the validated model can be utilized to generate new hypotheses. The hypotheses can be tested experimentally, providing another level of model validation. It is interesting to encounter cases when the experimental results do not match the model predictions, as this means that some aspects of the model should be reevaluated. Ultimately, this iterative process between mathematical modeling and experimental analysis improves our understanding of the underlying biology. The output of this step is a predictive model that can be used to better understand and optimize the biological system being studied.

## III. ILLUSTRATIVE EXAMPLES

Below, we present three examples of how we have implemented the process outlined above in our own research.

### A. LCK autoregulation

Lymphocyte-specific protein tyrosine kinase (LCK) mediates T cell activation. Activated T cells can release cytokines and cytotoxic factors to kill cancer cells. LCK’s activity is modulated via phosphorylation at two residues (Y394 and Y505). LCK catalyzes its own phosphorylation at both sites, and CSK, a regulatory kinase, phosphorylates LCK at the Y505 site. To better understand LCK regulation, we constructed a model composed of ODEs using mass action kinetics [9]. We followed the steps outlined above (sensitivity analysis, two-tiered parameter fitting, and comparison to validation data) to generate a predictive model of the dynamics and regulation of LCK phosphorylation.

We analyzed the optimal parameter sets, revealing statistically significant differences in LCK’s catalytic rates. These differences are dependent on LCK’s phosphorylation status. For instance, phosphorylation at Y394 increases LCK’s catalytic activity, while phosphorylation at Y505 reduced the catalytic activity. The eFAST analysis demonstrated that the association rate of LCK with itself or with CSK, and the dissociation and catalytic rates of CSK to the unphosphorylated form of LCK significantly influence the level of LCK phosphorylation. Interestingly, the model predicted novel autoregulatory feedback mechanisms that modulate LCK activity. Overall, our methodology not only estimated kinetic parameters of LCK phosphorylation, but also identified mechanisms involved in LCK regulation.

### B. TSP-1 and VEGF in breast cancer

We have also applied mathematical modeling to study angiogenesis, or the formation of new blood vessels. Tumors require their own blood supply to grow beyond the limits of oxygen and nutrient diffusion. Therefore, inhibiting angiogenic promoters, such as vascular endothelial growth factor (VEGF), and mimicking the action of inhibitors, such as thrombospondin-1 (TSP1), have a potential therapeutic role in limiting tumor development. We used the methods described here to develop a model of TSP1 and VEGF distribution in breast tumors [10]. We applied the model to help explain why some TSP1-derived anti-angiogenic drugs have not shown much success in the clinic. We also used the model parameter values and mechanistic predictions to hypothesize new ways to improve these therapies.

The sensitivity analysis of the model predicted that the concentration of the TSP1 receptor CD47 most significantly affects the main model output, the angiogenic ratio (the ratio of the pro- and anti-angiogenic signaling complexes). This quantity can be used to better understand why a TSP1 mimetic that targeted CD36 performed poorly in clinical trials. Furthermore, the model suggests that a CD47 binding mimetic may perform better. These predictions continue to be tested experimentally.

### C. NK cell signaling

As a final example, we present new results from our study to model Natural Killer (NK) cell activation leading to cytotoxic effects that target cancer cells. NK cell activation requires the integration of signaling inputs from multiple pathways [11]. A predictive computational model that quantifies the synergy between the multiple pathways involved in NK cell activation provides mechanistic insight into the NK cell’s killing potential and can accelerate the development of NK cell-based therapies.

We have constructed a molecularly-detailed model of three stimulatory pathways that contribute to overall NK cell activation: the CD16, NKG2D, and 2B4 pathways (**Fig. 2A**). These receptors activate MAPK, Akt, and PLCγ, signaling species that mediate NK cell activation and cytotoxicity. We produced a system of ODEs that incorporates biochemical reactions reported in the literature and predicts the signaling species’ concentrations. We trained the model using published experimental data (immunoblot measurements of key intracellular signaling species). Using the eFAST global sensitivity analyses, we first identified the most influential parameters that affect the main model outputs: the phosphorylated forms of the NK cell receptors, ERK, Akt, and PLCγ. We then applied PSO to estimate the values of the influential model parameters needed to match the training data. We validated the model using a separate set of data: measurements for NK cells simultaneously stimulated by multiple receptors or in the presence of molecular perturbations (e.g., RNAi, kinase inhibitors, or species knockdown). **Fig. 2B** shows predictions from the validated model compared to experimental data for ERK and Akt, following stimulation of the NKG2D receptor.

**Figure 2:**
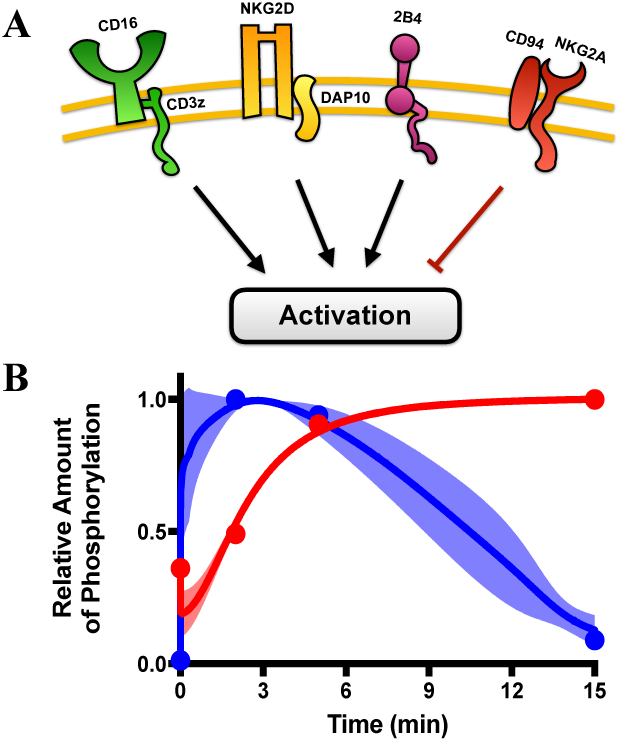
Relative level of phosphorylated ERK (red) and Akt (blue) upon NKG2D stimulation. *Lines*, mean model predictions. *Circles*, experimental data. *Shading*, 95% confidence interval.

Interestingly, although the CD16, 2B4, and NKG2D receptors all contribute to NK cell activation and cytotoxicity, their magnitude and mechanisms of action are quite different [11]. Our model shows that stimulation of CD16 produces a strong but transient response, compared to the other receptors. This can be explained by examining the estimated parameter values that characterize CD16 signaling: CD16 binds more tightly to its ligand and is more efficient in transducing the input signal (i.e., catalyzing subsequent reactions after ligand binding). Overall, the model provides mechanistic insight into the dynamics of different NK cell receptor stimulation and identifies ways to tune the signaling pathways for targeted cell-based therapies.

## ACKNOWLEDGMENT

The authors thank members of the Finley research group for helpful discussions and constructive feedback.

## REFERENCES

[1] M.P. Chapman and C.J. Tomlin, “Ordinary Differential Equations in Cancer Biology,” bioRxiv, available: https://doi.org/10.1101/071134

[2] K. Wilkie, P. Hahnfeldt, and L. Hlatky, “Using Ordinary Differential Equations to Explore Cancer-Immune Dynamics and Tumor Dormancy,” bioRxiv, available: https://doi.org/10.1101/049874

[3] J.R. Faeder, M.L. Blinov, and W.S. Hlavacek, “Rule-based Modeling of Biochemical Systems with BioNetGen,” Methods Mol. Biol., vol. 500, pp. 113–167, 2009.

[4] S. Marino, I.B. Hogue, C.J. Ray, and D.E. Kirschner, “A Methodology for Performing Global Uncertainty and Sensitivity Analysis in Systems Biology,” J. Theor. Biol., vol. 254, pp. 178–196, 2008.

[5] M.C.K. Khoo, Physiological Control Systems: Analysis, Simulation, and Estimation. Piscataway, NJ: IEEE Press, 2000.

[6] A. Raue, C. Kreutz, T. Maiwald, J. Bachmann, M. Scholling, U. Klingmuller, and J. Timmer. “Structural and Practical Identifiability Analysis of Partially Observed Dynamical Model by Exploiting the Profile Likelihood,” Bioinformatics, vol. 25, pp. 1923–1929, 2009.

[7] S.D. Finley, D Gupta, N Cheng, and D.J. Klinke, “Inferring Relevant Control Mechanisms for Interleukin-12 Signaling in Naïve CD4+ T cells,” Immunol. Cell Biol., vol. 89, pp. 100–110, 2011.

[8] S. Iadevaia, L.K. Nakhleh, R. Azencott, and P.T. Ram, “Mapping Network Motif Tunability and Robustness in the Design of Synthetic Signaling Circuits,” PLOS One, vol. 9, pp. e91743, 2014.

[9] J.A. Rohrs, P. Wang, and S.D. Finley, “Predictive Model of Lymphocyte-Specific Protein Tyrosine Kinase (LCK) Autoregulation,” Cell Mol. Bioeng, vol. 9, pp. 351–367, 2016.

[10] J.A. Rohrs, C.D. Sulistio, and S.D. Finley, “Predictive Model of Thrombospondin-1 and Vascular Endothelial Growth Factor in Breast Tumor Tissue,” npj Syst. Biol. Appl., vol. 2, article 16030, 2016.

[11] E.O. Long, H.S. Kim, D. Liu, M.E. Peterson, and S. Rajagopalan, “Controlling Natural Killer Cell Responses: Integration of Signals for Activation and Inhibition,” Annu. Rev. of Immunol., vol. 31, pp. 227–258, 2013.

